# Chlorogenic acid fails to confer neuroprotection in a chronic mouse model of Parkinson’s disease

**DOI:** 10.64898/2026.04.19.719432

**Authors:** Akshaya Rajan, Surya Prakash, Devendra Singh, Poonam Thakur

## Abstract

Parkinson’s disease (PD) is a progressive neurodegenerative disorder characterized by α-Synuclein (α-Syn) aggregation, dopaminergic neuronal loss, and chronic neuroinflammation. Chlorogenic acid (CA), a dietary polyphenol abundant in coffee, exhibits antioxidant and anti-inflammatory properties and has shown neuroprotective effects in acute toxin-based PD models. However, its efficacy in chronic, α-Syn–driven PD models remains unclear. Here, we evaluated the therapeutic potential of CA using an α-Syn–based *in vitro* system and a chronic α-Syn overexpression mouse model that recapitulates key pathological features of human PD.

*In vitro*, CA significantly improved cell viability, reduced α-Syn aggregation, and attenuated H_2_O_2_-induced apoptosis in U118 and N2a cells. In contrast, chronic oral administration of CA (100 mg/kg for 16 weeks) in C57BL/6J mice (male and female) failed to improve motor behavior, attenuate α-Syn pathology, preserve nigrostriatal dopaminergic neurons, or reduce oxidative stress–associated DNA double-strand breaks *in vivo*. Notably, CA elicited a modest reduction in microglial and astrocytic activation in female mice, highlighting a sex-dependent immunomodulatory response.

Collectively, these findings reveal a clear dissociation between robust *in vitro* neuroprotection and limited *in vivo* efficacy in a chronic α-Syn–driven PD mouse model, emphasizing the importance of incorporating progressive disease paradigms and sex as a biological variable in preclinical therapeutic evaluation.

## 1. Introduction

Parkinson’s disease (PD) is marked by progressive degeneration of dopaminergic neurons in the substantia nigra (SN) region of the brain [1], and aggregation of misfolded α-Synuclein (α-Syn) protein [2]. PD also exhibits pronounced sex differences in incidence, progression, symptom severity, and therapeutic response [3]. Despite substantial progress in understanding its molecular basis, PD remains incurable and continues to affect 2–3% of the elderly population worldwide [4]. Current treatments offer symptomatic management but fail to alter the disease progression, emphasizing the need for better neuroprotective or neurorestorative strategies capable of modifying the underlying pathology [5,6]. Dietary components with low toxicity and suitability for long-term administration have gained interest as potential disease-modifying agents [7,8]. Epidemiological studies report an inverse association between habitual coffee consumption and PD risk [9,10]. While caffeine has been extensively evaluated, its lack of efficacy in improving PD motor symptoms in randomized trials [11] suggests that other coffee-derived bioactive molecules may contribute to this protective association. Among these, Chlorogenic acid (CA) has emerged as a promising candidate due to its established antioxidant and anti-inflammatory properties [12].

CA is a common polyphenolic compound found in green coffee beans, tea, berries, cocoa, and several fruits [13,14]. Mechanistically, CA reduces oxidative stress by modulating poly (ADP-ribose) polymerase-1 (PARP1) [15], a key DNA repair enzyme implicated in parthanatos-mediated dopaminergic cell death [16,17]. CA suppresses PARP1 hyperactivation through RNF146 upregulation mediated by AKT1 signaling [18]. As elevated reactive oxygen species (ROS) and PAR formation accelerate α-Syn aggregation [19], modulation of the PARP axis represents a potential neuroprotective avenue. CA also mitigates rotenone-induced oxidative stress and improves behavioral outcomes in mice via PI3K/AKT-dependent mechanisms involving GSK-3β inactivation and GLP-1 release [20]. Beyond PD, CA demonstrates anti-inflammatory and antioxidative benefits across models of Huntington’s disease [21], Alzheimer’s disease [22], and metabolic dysfunction [23]. Given that PD patients often have dysregulated insulin signaling without explicit metabolic disorders [24,25]; CA can potentially restore these pathways to provide neuroprotection.

Preclinical evidence supports CA’s neuroprotective actions in toxin-based PD model systems. *In vitro*, CA enhances cell survival under 6-OHDA-induced stress and promotes autophagy [15]. *In vivo*, it reduces astrocytic activation and mitigates neurotoxin-induced pathology in MPTP and zebrafish models [26]. CA also exhibits favorable pharmacokinetics, with its predominant isomer, 5-caffeoylquinic acid, detectable in cerebrospinal fluid shortly after systemic administration [27,28]. However, all these studies have relied on acute toxin-induced PD models, that fail to reproduce key PD features such as α-Syn inclusion formation, dystrophic neurites, impaired axonal transport, and progressive neurodegeneration [29], contributing to the translational failure of many candidate neuroprotectants. Thus, CA’s relevance to disease-specific mechanisms of synucleinopathy using chronic, progressive, and α-Syn-driven PD models *in vitro* or *in vivo* remains unresolved.

Further, despite the well-recognized sex-based differences in PD pathogenesis, the majority of preclinical PD studies have historically relied on male animals, limiting the translational relevance of therapeutic findings [30]. Growing evidence indicates that neuroprotective and anti-inflammatory interventions may exert sex-specific effects, particularly in the context of α-synuclein pathology and glial activation [31]. Moreover, estrogen and estrogen-regulated signaling pathways have been shown to modulate oxidative stress, α-synuclein toxicity, PARP activity, and neuroinflammatory responses, all of which represent key mechanisms targeted by CA [32–34].

To address these limitations, the present study evaluates CA within a more pathophysiologically relevant framework using an α-Syn overexpression system combined with exogenous pre-formed fibrils (PFFs) in both *in vitro* and *in vivo* settings. This combinatorial model recapitulates hallmark PD features, including seeded α-Syn aggregation, chronic dopaminergic loss, and neuroinflammatory activation [35,36], thereby providing a stringent platform for assessing CA’s therapeutic potential in both sexes.

Our findings reveal that CA significantly improves cell viability and reduces α-Syn aggregate burden *in vitro*. In contrast, CA fails to rescue dopaminergic neurodegeneration or behavioral impairments in the chronic mouse model of PD, despite partially attenuating neuroinflammatory responses. Notably, this anti-inflammatory effect is more pronounced in female mice than in the males, identifying a sex-dependent therapeutic response not previously described for CA. Together, these results refine our understanding of CA’s context-dependent efficacy and emphasize the need for rigorous evaluation in chronic α-Syn-driven systems before advancing CA as a disease-modifying candidate for PD.

## 2. Materials and Methods

### 2.1. Cell culture

N2a and U118-MG cell lines were maintained in Dulbecco’s Modified Eagle Medium (DMEM; Hyclone) supplemented with 10% fetal bovine serum (FBS, Hyclone) and 1% Pen Strep (Gibco). Cells were cultured at 37°C in a humidified incubator under a 5% CO_2_ environment. For human α-Syn (h-α-Syn) overexpression, N2a and U118-MG cells were transduced with pLVX-Puro-h-α-Syn lentivirus and selected with puromycin (1 μg/ml for N2a; 4 μg/ml for U118-MG) and expression was verified using anti-h-α-Syn antibody (Table 1).

**Table 1.** List of antibodies used.

| Antigen | Catalogue Number | Host species | Dilution used |
| --- | --- | --- | --- |
| Tyrosine Hydroxylase | Abcam, ab112 | Rabbit | 1:600 |
| Tyrosine Hydroxylase | Abcam, ab76442 | Chicken | 1:700 |
| Alpha-synuclein (phospho S129) | Abcam, ab184674 | Mouse | 1:8000 |
| GFAP | Abcam, ab68428 | Rabbit | 1:1000 |
| Iba1 | Abcam, ab178847 | Rabbit | 1:1000 |
| Human-alpha-synuclein (Syn211) | Abcam, ab80627 | Mouse | 1:500 |
| Anti phospho H2AX-S139-3F2 | Abcam, ab22551 | Mouse | 1:750 |
| Recombinant human NRF2/NFE2L2 protein | Proteintech, ag9489 | Rabbit | 1:200 |
| PARP | Cell Signaling Technology, #9542s | Rabbit | 1:1000 |
| Anti-rabbit IgG H+L (Biotin) | Abcam, ab207995 | Goat | 1:500 |
| Anti-mouse IgG (H+L) AF488 | CST, #4408 | Goat | 1:1000 |
| Anti-chicken IgG (H+L) AF594 | Jackson Immuno Research, 703-585-155 | Donkey | 1:1000 |
| Anti-rabbit IgG (H+L) AF647 | CST, #4414 | Goat | 1:1000 |
| Anti-rabbit IgG HRP | Genei, 114038001A | Goat | 1:10000 |

### 2.2. Immunocytochemistry

U118-MG cells overexpressing h-α-Syn were plated on 13-mm glass coverslips placed in a 24-well culture plate for immunofluorescence studies. Cells were pretreated with CA for 12 hours, followed by exogenous addition of sonicated PFFs (30 seconds ON/30 seconds OFF, for a total of 20 minutes in a bath sonicator). After 9 hours of incubation with 100 nM PFFs, cells were fixed with 4% paraformaldehyde (PFA). After washing with 1X phosphate-buffered saline (PBS), the cells were incubated at room temperature (RT) in PBS containing 0.01% Triton X-100 (PBST), and 1% bovine serum albumin (BSA). Primary antibody (anti-h-α-Syn; Table 1) was added and incubated overnight at 4°C. Following PBST washes, cells were incubated with appropriate fluorophore-conjugated secondary antibody for one hour at RT. Cells were then incubated with 1 μM DAPI (HiMedia) and mounted using Vectashield Antifade Mounting Medium (Vector Labs).

### 2.3. MTT assay

N2a neuronal cells overexpressing h-α-Syn were seeded in a 96-well culture plate in triplicate for each condition. After a 12-hour pretreatment with CA, sonicated α-Syn PFFs were exogenously added to attain a final concentration of 25 nM. After 60 hours of PFF exposure, 0.5 mg/ml MTT [3-(4,5-Dimethylthiazol-2-yl)-2,5-Diphenyltetrazolium Bromide; ThermoFisher] was added to the cells and incubated for 4 hours at 37°C. The resulting MTT formazan crystals were solubilized using methanol, and absorbance was measured at 570 nm using a Spectramax-M5 plate reader. Average cell viability was calculated relative to the untreated control cells.

### 2.4. Western blotting

N2a cells were harvested 24 hours post-treatment with H_2_O_2_ (100 μM) and lysed in RIPA buffer (50 mM Tris–HCl, pH 7.4; 150 mM NaCl; 1 mM EDTA; 0.1% SDS; 0.25% sodium deoxycholate). The protein extracts were then centrifuged at 14000 rpm for 20 minutes at 4°C. The supernatant was collected, and the total protein concentration was determined using Bradford reagent (Bio-Rad). Proteins were separated by SDS–PAGE and transferred onto nitrocellulose membranes. Ponceau S (Sigma) staining was performed on the membranes after transfer to confirm equal protein loading and transfer efficiency. Membranes were imaged and then blocked with 5% skimmed milk in 1XTBST (20 mM Tris, 136 mM NaCl, pH 7.6 with 0.1% Tween-20) for an hour at room temperature. Membranes were incubated overnight at 4°C with anti-PARP antibody (Table 1). Following TBST washes, membranes were incubated with the HRP-conjugated anti-rabbit IgG secondary antibody for one hour and blots were developed using enhanced chemiluminescence (ECL) detection reagent and imaged on a ChemiDoc Imaging System (Bio-Rad).

### 2.5. Animal experimentation

An equal number of male and female ~12 weeks (W) old C57BL/6J mice were housed in the animal facility at IISER Thiruvananthapuram under a 12-hour light/dark cycle, with *ad libitum* access to food and water. All experimental procedures were approved by the Institutional Animal Ethics Committee (IAEC) in accordance with the guidelines established by the Committee for the Purpose of Control and Supervision of Experiments on Animals (CPCSEA), Government of India. Mice were randomly assigned to four experimental groups: (i) control (C), (ii) PD, (iii) PD+CA, and (iv) CA alone for both the sexes. To induce PD, a unilateral injection (right side of the mice brain) of the combination of rAAV-CMV-SYN-SNCA(WT)-WPRE-bGHpA 2/9 SNCA (Brain VTA, GT-0070; 0.5×10^10^ gc/site) and freshly sonicated PFFs (2.5 µg/site) in the medial SN (−3.2, −1.1, 4.6 for AP, ML and DV respectively) and the lateral SN (−3.2, −1.7, 4.15 for AP, ML and DV respectively) was performed. The control group was injected with an equivalent volume of 1X DPBS vehicle. Stereotactic surgeries were performed according to the procedure described earlier [37].

### 2.6. CA administration

CA (Sigma) was dissolved in PBS preheated to 40°C to obtain a final concentration of 20 mg/mL. It was administered once daily by oral gavage to the mice in the CA alone and PD+CA groups for 16 W, at a dose of 100 mg/kg body weight. Dosing was initiated one week post-surgery to allow for postoperative recovery and the initiation of PD pathogenesis. Vehicle (PBS) was orally administered to the C and PD groups in equivalent volumes. Body weight was recorded weekly throughout the study to monitor general health and potential treatment-related effects.

### 2.7. Pharmacokinetic analysis (LC-MS/MS)

Mice were administered 100 mg/kg CA by oral gavage, and body tissues, including blood plasma and brain, were collected at baseline (0 hour, pre-dosing) and at various time points post-dosing (0.5, 1, and 2 hours). CA concentrations in collected tissues were quantified externally by Aragen Life Sciences using liquid chromatography– tandem mass spectrometry (LC–MS/MS). Peak value was obtained in mice brain one hour post CA administration.

### 2.8. Behavioral tests

Behavioral tests were conducted at 8W and 16W post-surgery. Mice were allowed to acclimate for at least 30 min in a sound-attenuated behavioral testing room prior to testing. The open field, wire hang, corridor, and cylinder tests were performed at both time points as described previously [38]. The videos of the corridor test and cylinder test were analyzed by an experimenter blinded to the treatment groups.

### 2.9. Histology

All mice were euthanized by cervical dislocation 16W post CA administration. Transcardial perfusion was performed using chilled 1X DPBS followed by 4% PFA. The harvested brains were post-fixed in 4% PFA for 24 hours and subsequently stored in 20% sucrose. Coronal sections (25 μm) of SN and striatum (STR) were obtained using a vibratome (Leica VT1000 S). The sections were stored in PBS containing 0.1% sodium azide at 4°C. Free-floating sections were permeabilized with 0.1% Triton X-100 in PBS for 30 minutes at RT, followed by antigen retrieval in citrate buffer (pH 6.0) at 80°C for 30 minutes. Blocking was performed using 2% BSA in PBS with 0.1% Tween 20 for one hour at RT. The sections were incubated overnight at 4°C with primary antibodies (Table 1) prepared in PBS with 0.1% Tween-20. After washing, incubation with secondary antibodies (HRP-conjugated or AlexaFluor-conjugated) was carried out for 6 hours at room temperature or 4°C. Fluorescently stained sections were washed and mounted using Vectashield Antifade Mounting Medium (Vector Labs). For chromogenic staining, the sections were incubated with ABC reagent (Vectastain ABC Kit, Vector Labs) and developed using 0.05% Diaminobenzidine (Sigma) in 0.01% H_2_O_2_, followed by mounting with DPX (HiMedia) [36].

### 2.10. Imaging and quantification

Immunostained U118-MG cells were imaged at 63X magnification using an upright fluorescence microscope (Leica DM-6). Six fields per condition were captured, and the images were trained to detect h-α-Syn-specific puncta and differentiate them from the background using the Ilastik software. The resulting binary signal masks were quantified using the ImageJ particle analyzer.

Immunostained STR and SN sections were imaged at 4X and 10X magnification, respectively, using bright-field microscopy. Dopaminergic cell bodies in the SN were segmented from the background using Ilastik software. The masks generated were used to count the cells using the QuPath particle detector. Dopaminergic fiber density in the STR was assessed by measuring mean intensity using ImageJ. For quantification, 6–8 sections of SN and 8–10 sections of STR, spread across the rostro-caudal extent, from each mouse were used. Fluorescently stained sections were imaged at 20X magnification using a Leica upright fluorescence microscope, while 100X images were acquired with a Fluoview FV3000 inverted confocal microscope. For IBA1 (microglia), GFAP (astrocyte), and p-Syn (phosphorylated Serine-129 α-syn) analysis, 20X images of mid-SN slices from each brain were used to delineate the region of interest, and the percentage area occupied by the signal was calculated using ImageJ as shown previously [39]. For γH2AX quantification, three 100× images were captured along the medio–lateral extent of the SN per hemisphere from one representative section per animal across all experimental groups. The percentage area occupied by the signal was calculated after background subtraction and thresholding using ImageJ.

### 2.11. Statistical analysis

All data were plotted and analyzed using GraphPad Prism version 9.3.1. Data are expressed as mean ± standard error of the mean (SEM). Statistical comparisons were performed using one-way ANOVA followed by Tukey’s multiple comparisons post hoc test or two-way mixed-design ANOVA to examine effects between treatment groups and between hemispheres (injected vs. uninjected) within each group. When significant interaction effects were detected, simple effects were analyzed using Tukey’s multiple comparisons post hoc test. When interaction effects were not significant, main effects were interpreted independently, followed by Tukey’s multiple comparisons test for between-group analyses and Sidak’s multiple comparisons test for within-group (hemispheric) analyses. Statistical significance was set at p<0.05.

## 3. Results

### 3.1. CA did not improve PD-associated behavioral phenotypes and SN neurodegeneration

CA has previously been reported to improve cell survival under neurotoxic stress. Here, we assessed its neuroprotective efficacy in PD α-Syn overexpressing cells exposed to α-Syn PFF. As expected, exposure to PFF induced a robust increase in intracellular α-Syn aggregate levels (p<0.0001), which was significantly attenuated by CA treatment at both 10 μg/ml (p<0.001) and 100 μg/ml (p<0.0001) concentrations (Fig S1A and S1B). Consistent with these findings, PFF-induced reduction in cell viability (p<0.001) was rescued by CA at both 10 μg/ml (p<0.01) and 100 μg/ml (p<0.01) concentration (Fig S1C). Since CA is known to inhibit PARP accumulation and reduce apoptosis, PARP levels were assessed using immunoblotting (Fig S1D). CA modestly reduced full-length PARP and decreased H_2_O_2_-induced cleaved PARP levels, indicating reduced apoptosis. Overall, CA confers neuroprotection *in vitro* by attenuating PFF-induced α-Syn aggregation and reducing apoptosis, preserving cell viability under PD-relevant stress. We therefore sought to investigate the effects of CA in a chronic, PD-relevant *in vivo* model.

To investigate the effects of CA on PD progression and potential sex-dependent neuroprotection, equal numbers of male and female C57BL/6J mice were randomized into four experimental groups: (i) control (C), (ii) PD, (iii) PD+CA, and (iv) CA alone, followed by administration of specified treatments as depicted in Fig 1A. To confirm brain penetration of CA, LC-MS/MS analysis was performed one hour after oral dosing, detecting 1000±500 ng/mL in blood and 420±100 ng/g in brain tissue, thereby indicating effective blood–brain barrier permeability. Body weights remained stable across all groups throughout the experimental period, suggesting the absence of drug-related adverse effects during treatment (Fig S3).

**Fig 1.**
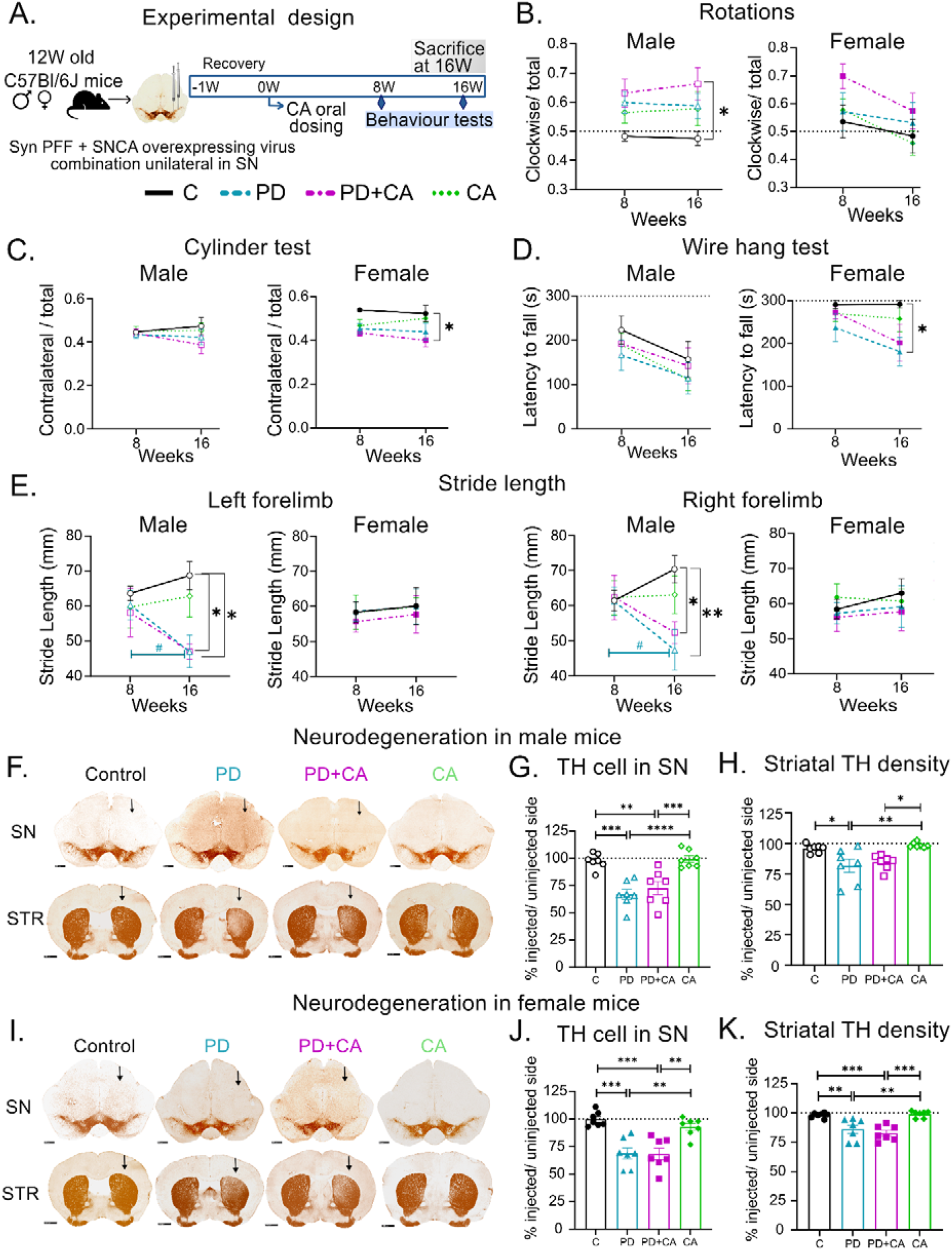
Lack of neuroprotection by CA in chronic mouse model of PD. (A) Schematic of the experimental timeline showing CA administration beginning one-week post-surgery, followed by behavioral assessments at 8W and 16W, and sacrifice at 16W. Injections were performed on the right hemisphere at medial and lateral SN. (B) Ratio of clockwise to total rotations in the open field test, showing a significant increase in male PD+CA mice compared to controls at 16W, indicative of unilateral lesioning. (C) Cylinder test analysis demonstrating reduced contralateral paw use in female PD+CA mice relative to controls at 16W. (D) Wire hang test showing greater grip strength loss in female PD mice at 16W compared to controls. (E) Stride length measurements from the corridor test showing significantly shortened left and right forelimb stride length in male PD and PD+CA groups relative to controls. (F) Representative TH-stained micrographs of the SN (scale bar: 500 µm) and STR (scale bar: 1000 µm) of male mice at 16W. (G) Quantification of TH-positive neurons in the SN demonstrates significant dopaminergic neurodegeneration on the injected side in PD and PD+CA males. (H) Striatal TH fiber density showing mild reductions in PD and PD+CA male groups. (I) Representative TH-stained micrographs of the SN (scale bar: 500 µm) and STR (scale bar: 1000 µm) of female mice at 16W. (J) Quantification of TH-positive neurons in the SN demonstrates significant dopaminergic neurodegeneration on the injected side in PD and PD+CA females. (K) Striatal TH fiber density showing mild reductions in PD and PD+CA female groups. All data are presented as mean±SEM; n = 7 mice per group per sex. Statistical analysis: repeated-measures two-way ANOVA for (B–E) and one-way ANOVA for (G, H, J, K), followed by Tukey’s or Sidak’s post hoc tests. Black arrow indicates injected side; *p<0.05, **p<0.01, ***p<0.001, ****p<0.0001, ^#^p<0.05. * indicates comparison between groups at a single timepoint; ^#^ indicates comparisons within group between 8W and 16W timepoints. Blue color ^#^ corresponds to PD-group.

Behavioral analysis was conducted at 8W and 16W post-surgery to determine whether CA mitigated PD-associated motor deficits. In the open field test, the ratio of clockwise rotations to total rotations was used as an index of unilateral lesion severity in mice brains (Fig 1B). Male PD+CA mice exhibited significantly increased clockwise rotations compared to control mice (p<0.05), while female PD+CA mice showed a similar, albeit non-significant, trend. No significant differences were observed in the remaining groups. Forelimb asymmetry resulting from unilateral lesions was further assessed using the cylinder test (Fig 1C). Male PD+CA mice showed reduced contralateral paw usage relative to controls. Similarly, female PD+CA mice exhibited significantly decreased contralateral paw usage at 16W compared to controls (p<0.05), indicating enhanced unilateral impairment. Similar to open field test, no other groups showed significant changes. Thus, motor asymmetry was more pronounced in PD+CA group as evaluated from these tests.

Muscular grip strength was evaluated using the wire hang test (Fig 1D). Male mice across all groups displayed a progressive decline in latency to fall over time but exhibited no significant intergroup differences. In contrast, female PD mice displayed a significant reduction in latency to fall at 16W compared to controls (p<0.05). The female PD+CA group also showed a similar trend toward reduced strength compared to controls, but this was not statistically significant.

Gait abnormalities were assessed by stride length analysis (Fig 1E). Male PD and PD+CA mice displayed significantly reduced left forelimb stride length (both p<0.05), along with reduced right forelimb stride length in PD (p<0.01) and PD+CA (p<0.05) groups relative to controls. Additionally, stride length in male PD mice declined significantly from 8W to 16W in both forelimbs (p<0.05). Interestingly, female treatment groups showed no significant stride length differences at 8W and 16W timepoints. Taken together, the persistence and progression of motor impairments across behavioral paradigms indicate that CA treatment failed to ameliorate any motor deficits observed in the PD groups in either sex.

All experimental groups were sacrificed post-16W, and their brains were harvested for histological evaluation. Overexpression of h-α-Syn in both PD and PD+CA groups was confirmed with a robust presence of Syn211 immunoreactivity (antibody specific for h-α-Syn) in both the SN and STR brain sections (Fig S4). Dopaminergic neuron integrity was then assessed using Tyrosine Hydroxylase (TH) immunostaining (Fig 1F–1K). Among males, the PD group showed a ~33% SN neuron loss (p<0.001), and the PD+CA group exhibited a ~27% loss (p<0.01) relative to controls (Fig 1G). In females, the PD and PD+CA groups showed ~31% (p<0.001) and ~32% (p<0.001) loss, respectively compared to controls (Fig 2J). Notably, the extent of neurodegeneration was comparable between PD and PD+CA groups in both sexes, indicating a lack of CA-mediated protection against neurodegeneration.

**Fig 2.**
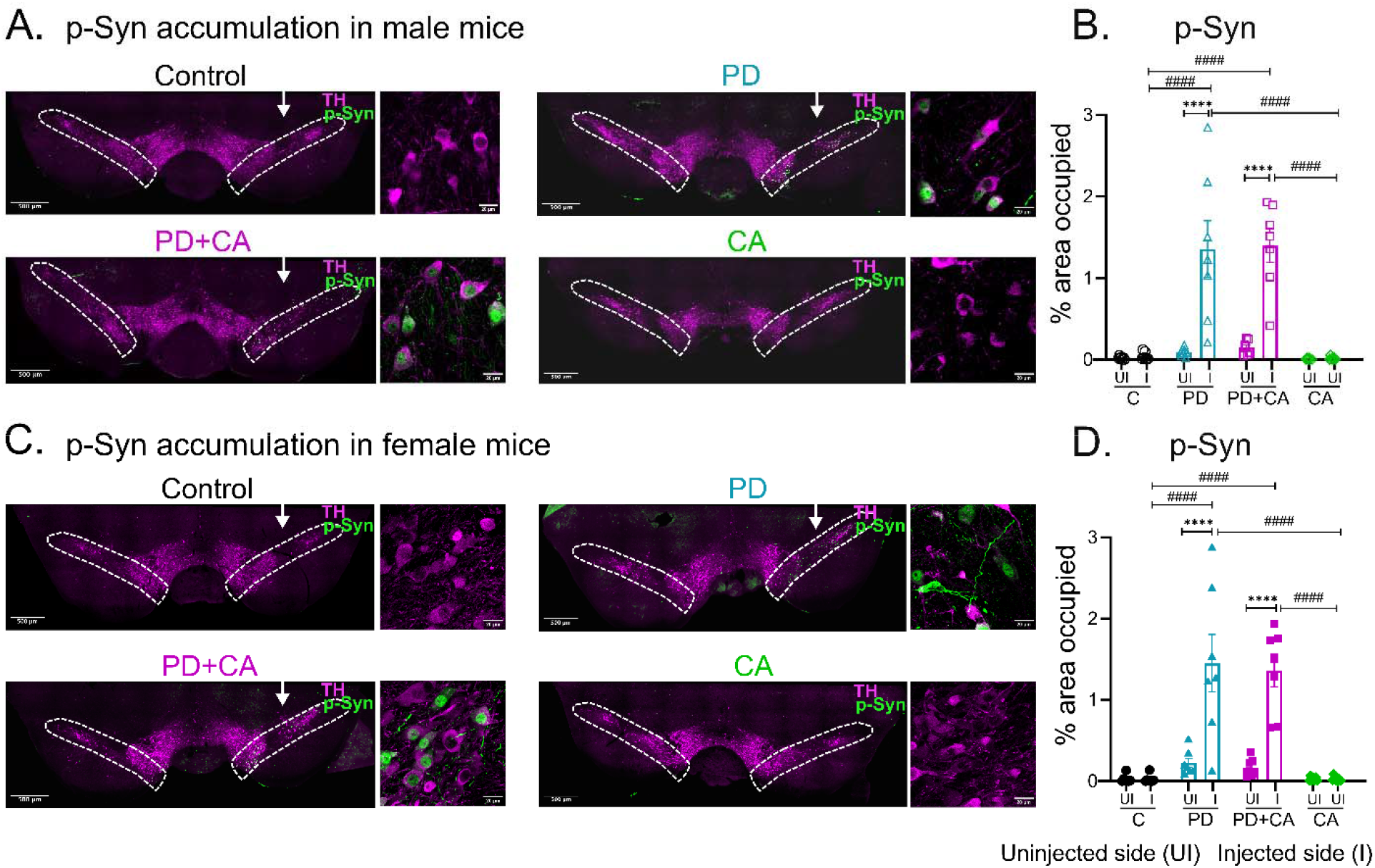
p-Syn accumulation in the SN is not reduced by CA treatment. (A) Representative TH and p-Syn co-stained images of the SN in male mice. (B) Quantification of p-Syn immunoreactivity in the SN (outlined by the white dotted line) of male mice. (C) Representative TH and p-Syn co-stained images of the SN in female mice, showing elevated p-Syn on the injected side in both PD and PD+CA groups. (D) Corresponding quantification of p-Syn from the SN region (outlined by the white dotted line) of female mice. White arrow indicates injected side. Scale bar: 500 μm for SN overview images; 20 μm for higher-magnification images of the injected side. Data are presented as mean±SEM; n = 7 per group for each sex. Statistical analysis was performed using mixed two-way ANOVA followed by Tukey’s multiple comparison test. ****p<0.0001; ^####^p<0.0001. * indicates comparisons within groups; # indicates comparisons between groups.

Similar trends were also observed for the STR TH density. Immunohistochemistry in STR revealed ~18% loss in male PD mice (p<0.05) and ~15% loss in PD+CA mice (p<0.05) compared to CA alone controls (Fig 1H). Female mice exhibited a similar pattern with the PD group showing ~14% loss (p<0.01), and the PD+CA group displaying ~17% loss (p<0.001) compared to controls (Fig 1K). As observed in the SN, striatal TH loss was comparable between PD and PD+CA groups in both sexes.

Collectively, these findings demonstrate that 16 weeks of oral CA administration neither mitigated dopaminergic neurodegeneration, nor improved behavioral deficits in this progressive α-Syn–based PD mouse model, indicating a lack of neuroprotective efficacy *in vivo*.

### 3.2. p-Syn deposition remained unchanged following CA administration

Aggregation and deposition of α-Syn fibrils, culminating in Lewy body formation, are hallmark pathological features of human PD. To assess α-Syn aggregation, phosphorylated Serine-129 α-Syn (p-Syn) immunoreactivity was quantified in SN. In male mice, both PD and PD+CA groups showed significantly higher p-Syn levels in their injected side compared to the uninjected side (p<0.0001) (Fig 2A and 2B). These groups also had higher p-Syn deposition than that observed in the injected side of the control group (p<0.0001). Similarly, in females, the PD and PD+CA groups showed significantly higher p-Syn accumulation in their injected side relative to their uninjected side (p<0.0001) and compared to the injected side of the control group (p<0.0001) (Fig 2C and 2D). Moreover, p-Syn accumulation in SN remained similar in chronic PD male and female mice at 16W. Despite sustained CA exposure, PD-associated p-Syn accumulation in the SN remained unchanged in both sexes after 16 weeks, indicating a lack of effect on α-synuclein pathology.

### 3.3. CA attenuated neuroinflammation in female but not male mice

In addition to dopaminergic neurodegeneration and p-Syn accumulation in the SN, neuroinflammation is a key contributor to PD pathogenesis. Neuroinflammatory responses were assessed by quantifying microglial activation as the percentage area occupied by IBA1-positive cells in the SN. In males, IBA1 levels were significantly increased on the injected side in both the PD group (1.5 fold; p<0.01) and the PD+CA group (1.6 fold; p<0.01) relative to their respective uninjected sides (Fig 3A and 3B). IBA1 levels were also significantly higher than those on the sham-injected side of control mice in both PD (1.8 fold; p<0.01) and PD+CA (1.7 fold; p<0.01) groups. Importantly, no significant difference was seen between the injected sides of PD and PD+CA male mice, indicating that CA did not reduce microglial activation in males.

**Fig 3.**
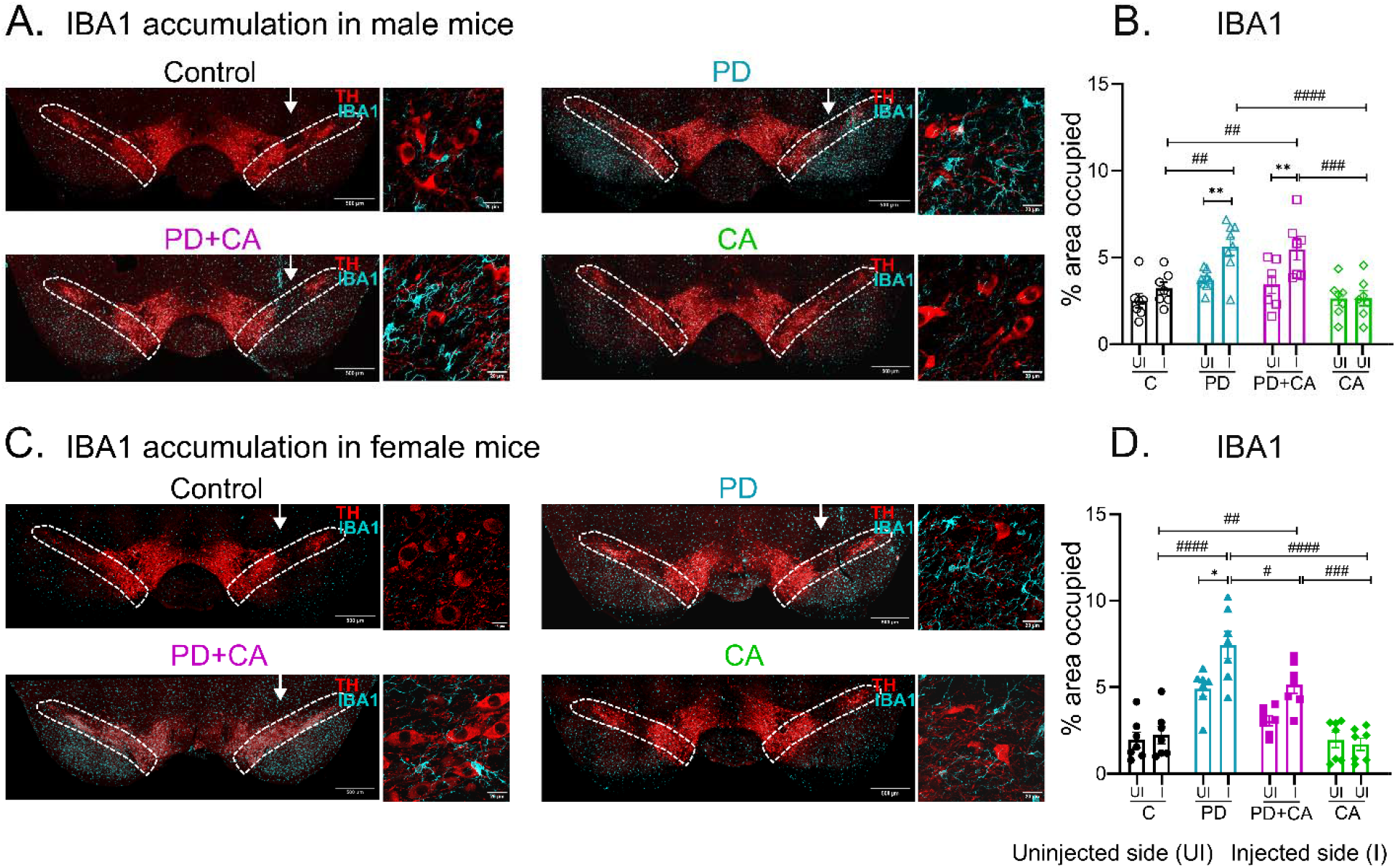
CA shows mild attenuation of microglial activation in female mice in a chronic PD model. (A) Representative images of SN co-stained with TH and IBA1 in male mice showing similar IBA1 accumulation in PD and PD+CA groups. (B) Quantification of IBA1 in SN (outlined by white dotted line) of male mice demonstrates no significant effect of CA treatment. (C) Representative images of SN co-stained with TH and IBA1 in female mice, indicating reduced microglial accumulation in the PD+CA group relative to PD. (D) Quantification of IBA1-positive area in the SN (outlined by white dotted line) in females. White arrow indicates injected side. Scale bar-500 μm for the SN overview image; 20 μm for higher-magnification images of the injected side. Data represented as mean±SEM; n=7-8 per group for each sex; Statistical analysis was performed using mixed two-way ANOVA followed by Tukey’s or Sidak’s multiple comparison test as post-hoc analysis; *p<0.05, **p<0.01, ^#^p<0.05 ^##^p<0.01, ^###^p<0.001, ^####^p<0.0001. * indicates comparisons within groups; # indicates comparisons between groups.

In contrast, females exhibited a differential response. IBA1 staining was significantly higher on the injected side in both PD (1.5 fold; p<0.05) and non-significantly higher in PD+CA (1.7 fold; p>0.05) groups compared to their uninjected sides (Fig 3C and 3D). IBA1 expression was also markedly higher on the injected side of PD (3.3 fold; p<0.0001) and PD+CA (2.3 fold; p<0.01) female mice compared to the injected side of controls. Notably, in females, IBA1 accumulation on the injected side was significantly reduced in the PD+CA group compared with the PD group (p<0.05), indicating CA-mediated attenuation of microglial activation in females.

Apart from microglia, astrocytic activation, assessed by GFAP immunostaining, known to increase during PD progression to aid in clearing protein aggregates and cellular debris [40], was also observed. In male mice, GFAP levels on the injected side were ~3 fold higher in the PD group (p<0.05) and ~4.5 fold higher in the PD+CA group (p<0.001) compared to their respective uninjected sides (Fig 4A and 4B). In female mice, GFAP levels were elevated by 3.2 fold in the PD group (p<0.0001) and 2.5 fold in the PD+CA group (p<0.05) on the injected side relative to their uninjected sides (Fig 4C and 4D). Notably, GFAP accumulation was significantly lower in females from the PD+CA group compared with the PD group (p<0.05).

**Fig 4.**
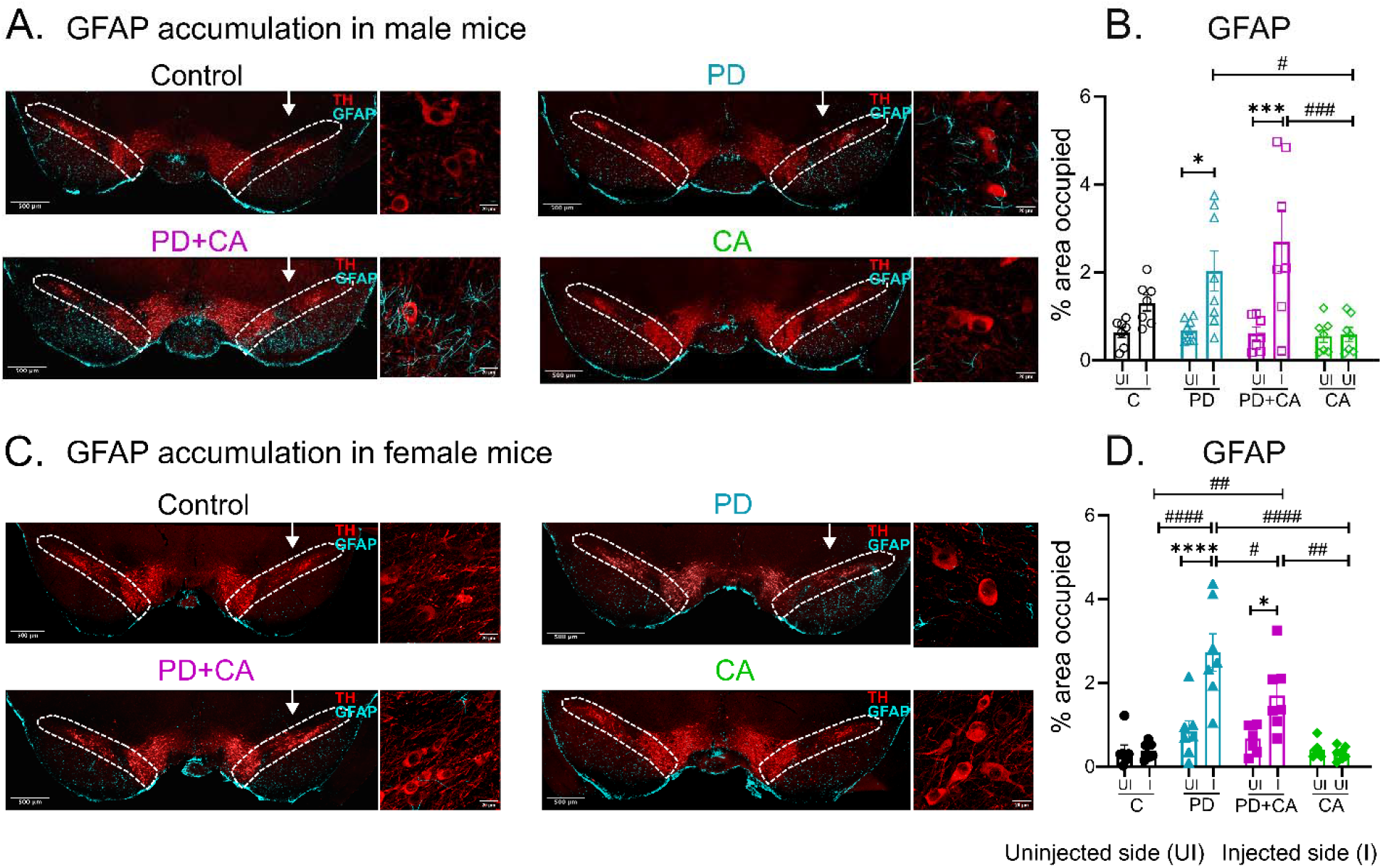
CA selectively reduces astrocytic reactivity in female mice in a chronic PD model. (A) Representative images for SN co-stained with TH and GFAP in male mice showing similar GFAP with and without CA in the PD groups. (B) Quantification of GFAP-positive area in SN (outlined by white dotted line) of male mice. (C) Representative images for SN co-stained with TH and GFAP in female mice, indicating reduced GFAP accumulation with CA. (D) Quantification of GFAP-positive area in SN (outlined by white dotted line) of female mice. White arrows indicate injected side. Scale bar-500 μm for SN overview image and 20 μm for higher-magnification images of the injected side. Data represented as mean±SEM; n=7-8 per group for each sex; Statistical analysis was performed using mixed two-way ANOVA followed by Tukey’s multiple comparison test as post-hoc analysis; *p<0.05, ***p<0.001, ****p<0.0001, ^#^p<0.05, ^##^p<0.01, ^###^p<0.001, ^####^p<0.0001, * indicates comparison within groups, and ^#^ represents comparison between groups.

Together, the reductions in both IBA1 and GFAP immunoreactivity in female PD+CA mice indicate a modest but sex-specific anti-inflammatory effect of CA. While neuroinflammatory markers were elevated in both male and female PD mice, CA treatment selectively attenuated astrocytic and microglial activation in females, with no detectable anti-inflammatory effect in males, highlighting a sex-dependent therapeutic response.

### 3.4 PD-associated oxidative stress is not attenuated by CA in the chronic mouse model

Oxidative stress is a major contributor to PD pathology, arising from an imbalance between reactive oxygen species (ROS) and antioxidant defenses that ultimately promotes dopaminergic neurodegeneration. Excessive ROS can induce DNA double-strand breaks (DSBs). Therefore, phosphorylated histone H2AX (γH2AX), a well-established marker of DNA damage, was used as a readout of oxidative stress in the SN across experimental groups. CA has previously been reported to reduce oxidative stress and associated DNA damage in neurotoxin-based PD models [41,42].

In male mice, γH2AX levels on the injected side were increased by ~6.8 fold in the PD group (p<0.01) and ~7.8 fold in the PD+CA group (p<0.05) relative to their respective uninjected sides (Fig 5A and 5B). γH2AX accumulation was also significantly higher in the PD (5.3 fold; p<0.01) and non-significantly higher in the PD+CA (3.7 fold; p>0.05) groups compared with the injected side of control mice. Similarly, in female mice, γH2AX accumulation on the injected side increased by ~3.9 fold in the PD group (p<0.05) and ~4 fold in the PD+CA group (p<0.001) relative to their uninjected sides (Fig 5C and 5D). Compared with the sham-injected side of control female mice group, γH2AX levels were significantly elevated in both PD (4.5 fold; p<0.05) and PD+CA (5.7 fold; p<0.001) groups. Notably, γH2AX accumulation was comparable between PD and PD+CA groups in both sexes, indicating that CA treatment for 16 weeks did not attenuate DSBs accumulation or oxidative stress in this chronic PD mice model. These findings suggest that the antioxidative effects of CA previously observed in acute or toxin-based PD models do not translate to progressive α-Syn–driven pathology.

**Fig 5.**
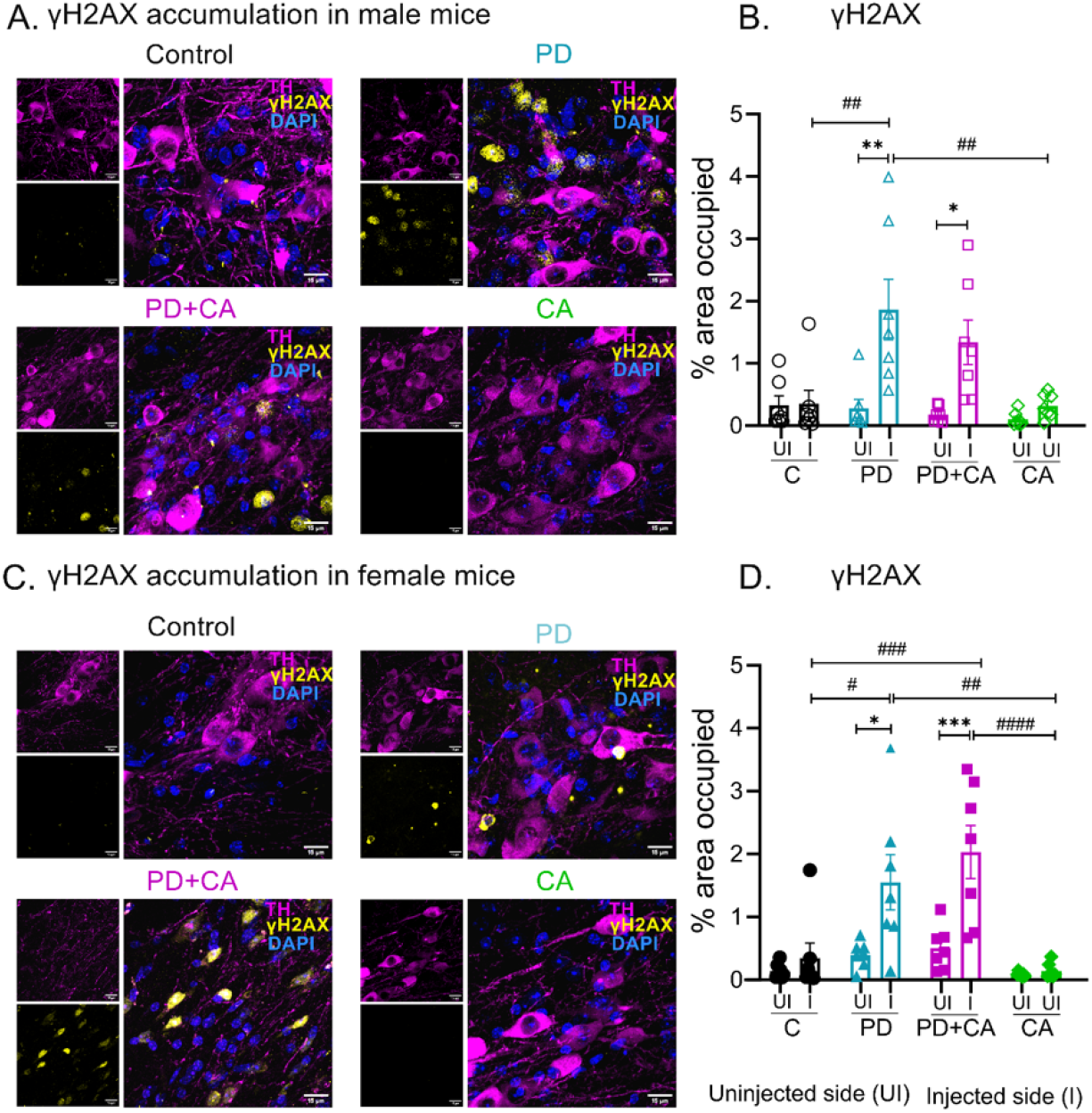
DNA damage is increased in chronic PD and is not alleviated by CA treatment. (A) Representative γH2AX and TH co-stained images of the injected SN across different experimental groups in male mice. (B) Quantification of γH2AX immunoreactivity in male mice demonstrating elevated DSBs on the injected side in PD and PD+CA male mice groups. (C) Representative γH2AX and TH co-stained images of the injected SN across indicated experimental groups in female mice. (D) Quantification of γH2AX immunoreactivity in female mice showing increased DSBs accumulation on the injected side in PD and PD+CA groups. Scale bar-15 μm; Data represented as mean±SEM; n=7-8 per group for each sex; Statistical analysis was performed using mixed two-way ANOVA followed by Tukey’s multiple comparison test as post-hoc analysis; *p<0.05, **p<0.01, ***p<0.001, ^#^p<0.05, ^##^p<0.01, ^###^p<0.001, ^####^p<0.0001, * indicates comparison within groups, and ^#^ indicates comparison between different groups.

Consistent with these observations, γH2AX immunoreactivity was also detected in the STR, the terminal region of dopaminergic neurons (Fig S5). Furthermore, an additional marker associated with antioxidant response, NRF2 (Nuclear factor erythroid 2-related factor 2) showed comparable expression levels in the STR of PD and PD+CA groups (Fig S6), further supporting the lack of measurable CA-mediated modulation of oxidative stress *in vivo*.

## 4. Discussion

In this study, we comprehensively evaluated the neurotherapeutic potential of CA -a dietary polyphenol with a well-established safety profile-in cellular and *in vivo* mice models of PD based on α-Syn pathology. Previous studies have utilized diverse systems such as H_2_O_2_ and 6-OHDA to assess CA’s modulatory effects [15,18,26]. Here, we provide the first evidence of its efficacy in a neuronal model combining human α-Syn overexpression with exogenous PFF exposure, a “dual-hit” system that robustly recapitulates seeded α-Syn aggregation, a pathological feature typically absent in simple α-Syn overexpression models. Consistent with previous cellular PD studies, CA significantly enhanced neuronal viability, reduced α-Syn aggregation, and decreased PARP cleavage indicating suppression of apoptotic signaling. These *in vitro* findings align with CA’s reported ability to mitigate oxidative stress, modulate PARP activity, and alleviate proteotoxic stress under acute pathological insults. However, our *in vivo* results demonstrate that these protective effects did not translate to sustained benefit in the setting of chronic α-Syn overexpression and protracted neurodegenerative stress.

The reported LD_50_ of CA is approximately 1,580 mg/kg[43]. Following standard pharmacological practice, we selected an *in vivo* dose corresponding to approximately 5-10% of the LD_50_, a range commonly used to define the upper limit of tolerability for repeated dosing. Consistent with this approach, previous study using acute PD models have reported dose-dependent motor improvement and neuroprotection effects of CA, with maximum efficacy observed at 100 mg/kg administered for three weeks [44]. Similarly, in a 6-OHDA rat model, CA administered at the dose of 60 mg/kg for 7 days improved motor performance and striatal dopamine levels, accompanied by reduced α-Syn expression and attenuated apoptosis, although dopaminergic cell rescue was not quantified [45]. Moreover, higher doses were deliberately avoided in the present study, as our *in vitro* experiments showed clear cytotoxicity at elevated concentrations (>150 ug/ml), raising concerns about potential toxicity *in vivo*. Accordingly, CA was administered at 100 mg/kg, a dose that is among the highest reported in the literature and represents the safe and pharmacologically justified dose for long-term administration. The absence of mortality and maintenance of comparable body weights between CA-treated and vehicle-treated mice confirms absence of any drug-induced toxicity *in vivo*. In addition, pharmacokinetic analysis by LC-MS/MS confirmed that CA reached biologically relevant concentrations in the brain.

Despite confirmed brain exposure and prolonged administration, CA failed to confer neuroprotection in our chronic PD model. Chronic oral CA administration of 16 weeks failed to preserve nigral dopaminergic neuron survival or striatal dopaminergic terminal integrity. CA treatment also did not reduce the accumulation of p-Syn within the SN, suggesting its inability to disrupt the propagation or facilitate the clearance of pathogenic α-Syn species. Given that p-Syn is a cardinal feature of Lewy pathology and closely linked to neurotoxicity, its persistence likely contributes to the unabated dopaminergic degeneration observed. These results suggest that CA does not likely engage α-Syn-specific clearance mechanisms, such as the autophagy-lysosomal pathway or the ubiquitin-proteasome system, under chronic disease conditions.

Biological sex is increasingly recognized as a critical modifier of neurodegenerative trajectories and therapeutic responsiveness in PD, yet it remains underrepresented in preclinical disease modeling. Sex-dependent differences in neuroimmune regulation and cellular stress responses may critically influence drug efficacy under chronic pathological conditions. By evaluating CA in both male and female mice, our study enables the identification of sex-dependent effects that would likely be missed in single-sex study designs, strengthening the translational relevance of the findings. Interestingly, CA elicited a modest yet significant anti-inflammatory response in a sex-dependent manner. In female mice, CA treatment significantly suppressed microglial and astrocytic activation in the SN, as reflected by reduced immunoreactivity for IBA1 and GFAP, respectively. No such immunomodulatory effect was detected in male mice, despite comparable levels of α-Syn pathology and neuronal loss. This sexual dichotomy may be rooted in established differences in neuroimmune regulation, potentially involving estrogen signaling. Estrogen is known to exert anti-inflammatory and neuroprotective effects and can promote expression of RNF146, an E3 ubiquitin ligase involved in PAR catabolism [18,34]. A plausible convergence between estrogen-mediated pathways and CA’s reported PARP-inhibitory activity may thus underlie the enhanced immunomodulatory response observed in females. Nonetheless, this attenuation of neuroinflammation did not confer neuroprotection, indicating that inflammatory modulation alone is insufficient to arrest neurodegeneration once chronic pathology is established.

Oxidative stress-induced DNA damage constitutes a pivotal mechanism of neuronal vulnerability in PD [46]. Elevated γH2AX levels have been associated with increased α-Syn aggregation and propagation, processes that are critical drivers of PD pathology [47]. Significantly high genomic oxidative damage activates PARP that can further promote neurodegeneration[48]. As γH2AX represents an associative marker of oxidative DNA damage linked to PARP activation, its evaluation enables meaningful comparison of CA–mediated effects in a chronic PD model [49]. Assessment of γH2AX, a marker of DNA DSBs, revealed substantial genomic damage in the SN of both male and female PD mice, which was not ameliorated by CA. The persistence of the γH2AX signal despite prolonged CA administration suggests that CA cannot counteract the relentless oxidative stress or associated DNA damage once a chronic neurodegenerative state is established. Although CA is a well-characterized NRF2 activator and antioxidant, elevated levels of γH2AX persisted in the nigrostriatal pathway of CA-treated PD mice. Furthermore, another oxidative stress-related marker, NRF2, showed no improvement with CA treatment.

These collective findings indicate that CA is ineffective against the multifactorial and sustained oxidative burden inherent to chronic α-Syn-driven pathology. In such setting, DNA damage likely arises not only from ROS but also from concurrent mitochondrial dysfunction, calcium dyshomeostasis, ferroptosis-linked lipid peroxidation, and paracrine inflammatory signals[46]. Consequently, therapeutic strategies targeting a single antioxidant pathway may exhibit limited efficacy in advanced or chronic PD models where oxidative injury accrues via a convergent network of mechanisms.

The equivalent magnitude of dopaminergic neuron loss in PD and PD+CA cohorts of both sexes conclusively demonstrates that CA does not alter the core neurodegenerative trajectory in this synucleinopathy model. This outcome stands in stark contrast to numerous reports of CA-mediated neuroprotection in acute toxin-based PD models (e.g., MPTP, 6-OHDA), where oxidative stress and mitochondrial dysfunction are primary pathogenic drivers. In such acute models, antioxidant and anti-apoptotic interventions frequently yield robust neuroprotection. By contrast, the model employed in our study, which is produced by adenoviral α-Syn overexpression along with PFF co-injection, more closely reflects the progressive, proteinopathy-centric nature of human PD, suggesting that CA’s pharmacological actions may be better suited to counteracting acute oxidative insults rather than chronic proteostatic failure.

CA’s inability to modulate α-Syn propagation, prevent neurodegeneration, or mitigate oxidative genomic damage suggests that it is unlikely to be effective as a monotherapy for established PD. Nevertheless, its sex-specific anti-inflammatory activity *in vivo* and robust cytoprotective effects *in vitro* suggest potential utility under alternative therapeutic contexts. In our experimental design, treatment was initiated one week after pathology induction to mirror a clinical scenario where therapy begins after diagnosis. However, CA may hold greater promise as a preventive or early-intervention agent when oxidative and inflammatory pathways remain more amenable to pharmacological modulation. Furthermore, our results support the view that effective disease modification in PD will likely require combinatorial approaches that directly target α-Syn aggregation, enhance proteostatic clearance mechanisms, and provide sustained antioxidant support.

Importantly, the lack of efficacy observed with this dosing regimen—previously shown to be neuroprotective in acute PD models—is unlikely to result from insufficient dosing or inadequate brain bioavailability. Instead, these findings indicate that CA efficacy may depend on disease stage or treatment timing, and that pre-treatment or alternative dosing strategies may be required to counter progressive neurodegeneration. More broadly, our results underscore the importance of evaluating candidate therapeutics in preclinical systems that closely emulate the protracted pathology of human PD. The slow progression of PD is often poorly modeled by acute neurotoxicant paradigms, and several interventions that demonstrated robust neuroprotection in acute models, including certain α-Syn immunotherapies and GLP-1 receptor agonists, have subsequently failed to meet primary endpoints in clinical trials [50,51].

In summary, although CA potently reduced α-Syn aggregation and neuronal death in PFF-challenged cellular models, these beneficial effects did not translate into meaningful neuroprotection in a progressive α-Syn-driven mouse model. Despite confirmed brain bioavailability and prolonged administration, CA failed to ameliorate behavioral deficits, dopaminergic neurodegeneration, pathological p-Syn accumulation, or oxidative DNA damage *in vivo*. More broadly, our study emphasizes careful consideration of model choice in translational neurodegeneration research and highlight the limitations of relying on acute paradigms for predicting clinical efficacy. By systematically incorporating sex as a biological variable, this study further underlines the importance of biological context in clinically relevant assessment of potential therapeutic candidate for PD.

## Abbreviations

PD: Parkinson’s disease)
α-Syn: Alpha-Synuclein)
CA: Chlorogenic acid)
N2a: Neuro-2A)
MTT: 3-(4,5-Dimethylthiazol-2-yl)-2,5-diphenyltetrazolium bromide)
PARP-1: Poly ADP-Ribose Polymerase-1)
PFF: Pre-formed fibrils of α-synuclein)
STR: Striatum)
SN: Substantia nigra)
TH: Tyrosine hydroxylase)
p-Syn: Serine-129 phosphorylated α-synuclein)
IBA1: Ionized calcium binding adaptor molecule 1)
GFAP: Glial fibrillary acidic protein)
γH2AX: Serine-139 phosphorylated histone variant H2AX)
NRF2: Nuclear factor erythroid 2-related factor 2

## Data Availability

The data are available from the corresponding author on request.

## Acknowledgements

We thank Santhosh Kumar Subramanya and Unnati Agrawal for helping with the animal experiments. We also thank Patange Pavan Eknath for help with the behavioral analysis.

## Author Information

### Authors and affiliation

School of Biology, Indian Institute of Science Education and Research, Thiruvananthapuram, Kerala, 695551, India Akshaya Rajan, Surya Prakash, Devendra Singh, Poonam Thakur

### Contributions

Conceptualization: Poonam Thakur; Methodology: Akshaya Rajan, Surya Prakash, Devendra Singh; Formal analysis and investigation: Akshaya Rajan and Devendra Singh; Writing - original draft preparation: Akshaya Rajan and Surya Prakash; Writing - review and editing: Akshaya Rajan, Devendra Singh and Poonam Thakur; Funding acquisition: Poonam Thakur; Resources: Poonam Thakur; Supervision: Poonam Thakur

## Ethics declarations

### Ethical approval

All experimental procedures were approved by the Institutional Animal Ethics Committee (IAEC) in accordance with the guidelines established by the Committee for the Purpose of Control and Supervision of Experiments on Animals (CPCSEA), Government of India.

