## Supplementary material for "Chlorogenic acid fails to confer neuroprotection in a chronic mouse model of Parkinson’s disease"

**Supplementary Information**

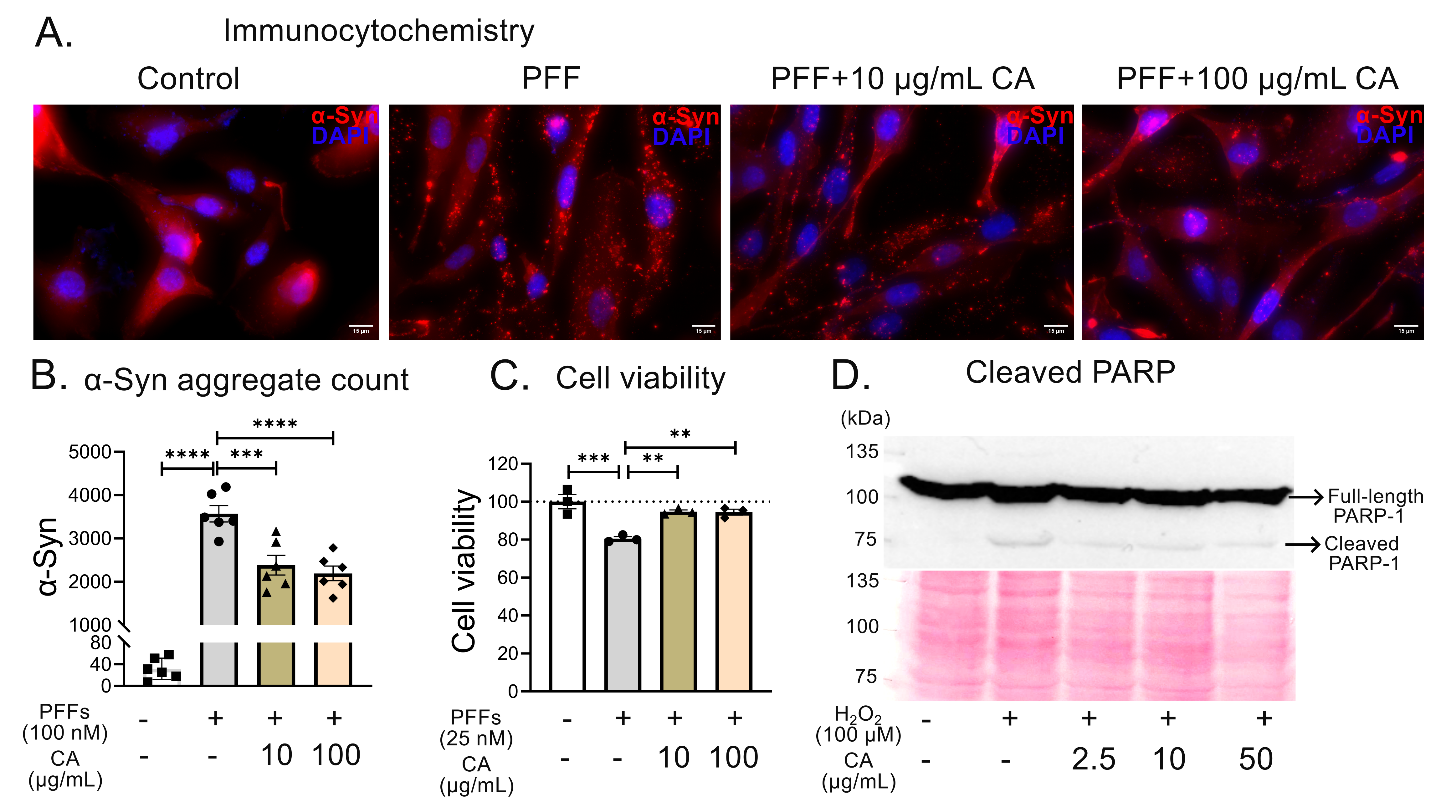

**Fig S1- CA improved cell survival and reduced α-Syn accumulation in PFF-treated h-α-Syn overexpressing cells** *(A) Representative micrographs of U118-MG cells immunostained for h-α-Syn and counterstained with DAPI (Scale bar- 15 μm). (B) Quantification of h-α-Syn-positive puncta shows reduced α-Syn accumulation following co-incubation with 100 nM PFFs and CA. (C) MTT assay demonstrates improved N2a cell viability after 12 hours of CA pre-treatment followed by 60 hours of co-incubation with 25 nM PFFs and CA. (D) Representative immunoblot of full-length PARP-1 and cleaved PARP showing decreased apoptosis with CA treatment (uncropped blot provided in Fig S2). Data presented as mean ± SEM, ordinary one-way ANOVA followed by Tukey’s multiple comparison test as post-hoc analysis; n=6 per group in B and 3 per group in C; **p<0.01, ***p<0.001, ****p<0.0001.*

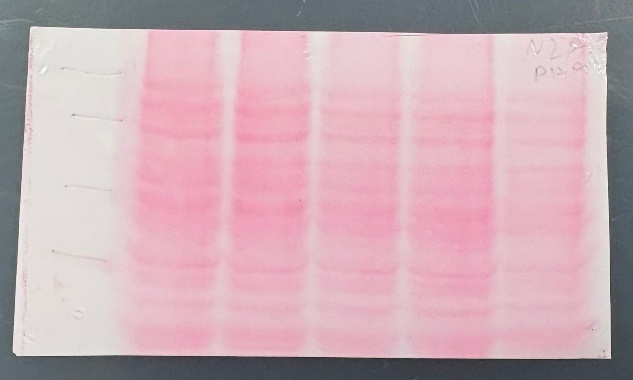

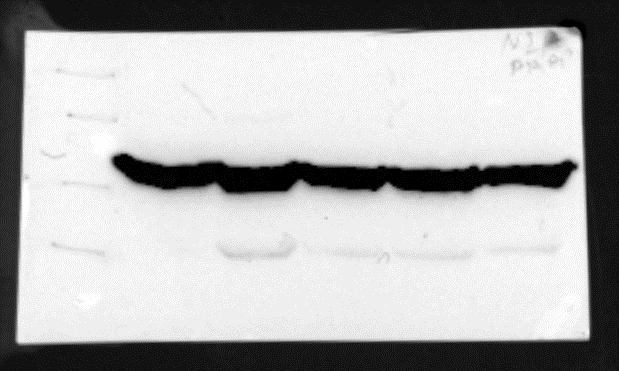

PARP-1 molecular weight - 116kDa; Cleaved PARP-1 molecular weight – 89kDa

Ladder	control	CA0	CA2.5	CA10	CA50

100 μM H_2_O_2_ (24hours)

180

135

100

75

kDa

Ladder	control	CA0	CA2.5	CA10	CA50

180

135

100

75

kDa

100 μM H_2_O_2_ (24hours)

**Fig S2-** Full sized or uncropped blot stained for ponceau and probed for PARP-1 with molecular ladders (marked with pencil) corresponding to the Fig S1D. The dotted lines denote the region shown in Fig S1D.

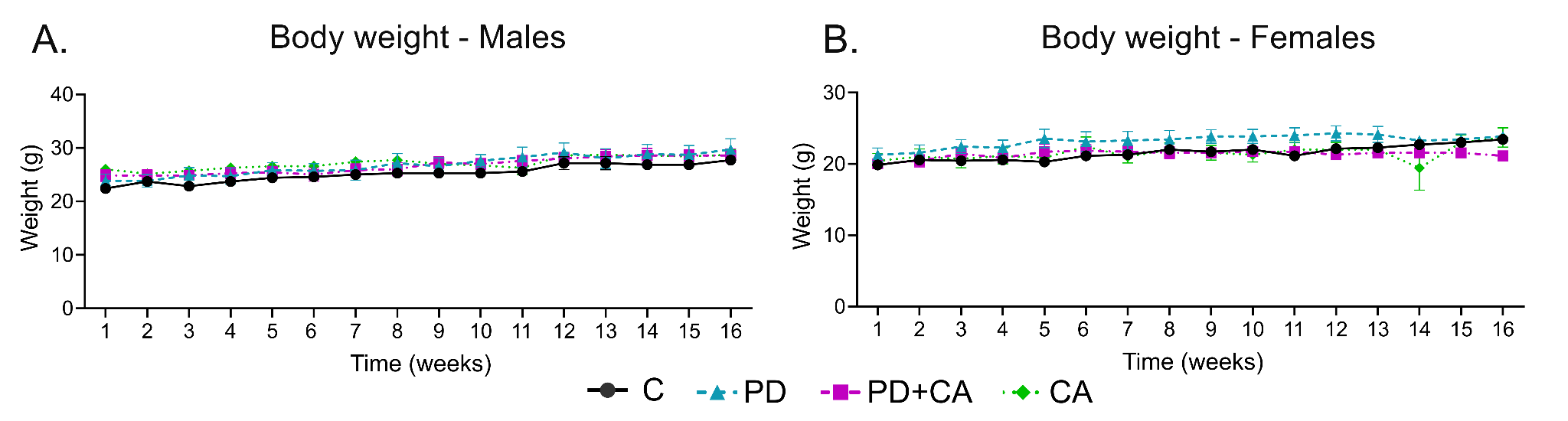

**Fig S3- Body weight remains unchanged across experimental groups.** Body weights of A. male and B. female mice recorded longitudinally throughout the study period. Data are presented as mean ± SEM; n = 7 mice per group.

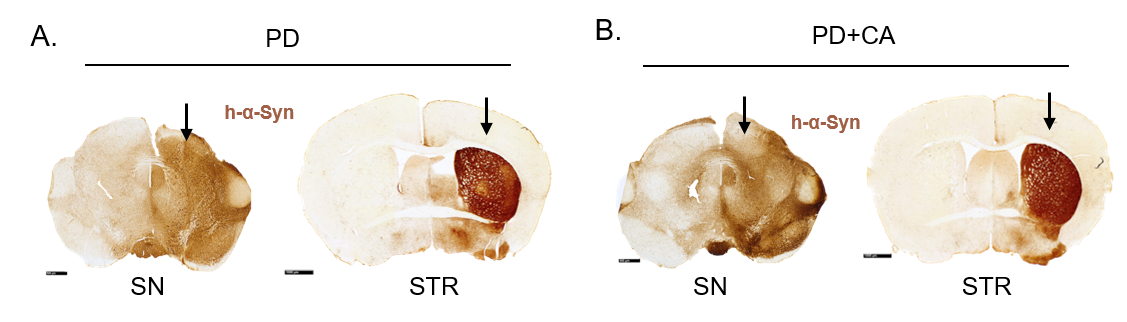

**Fig S4- Confirmation of h-α-synuclein overexpression in PD groups.** Representative images showing human α-synuclein (h-α-Syn; Syn211 antibody) immunoreactivity in the SN and STR of virus-injected mice from the A. PD and B. PD+CA groups. Scale bars: 500 μm (SN) and 1000 μm (STR); black arrow indicates the injected side.

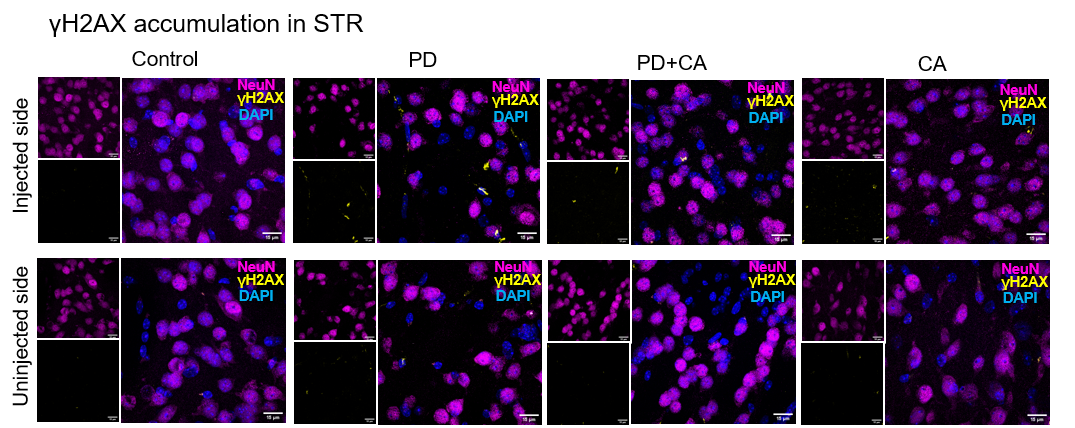

**Fig S5- γH2AX accumulation in the striatum (STR).** Representative images of the STR co-stained for γH2AX (yellow), NeuN (magenta), and DAPI (blue) in the injected and uninjected hemispheres across all experimental groups. Scale bar: 15 μm

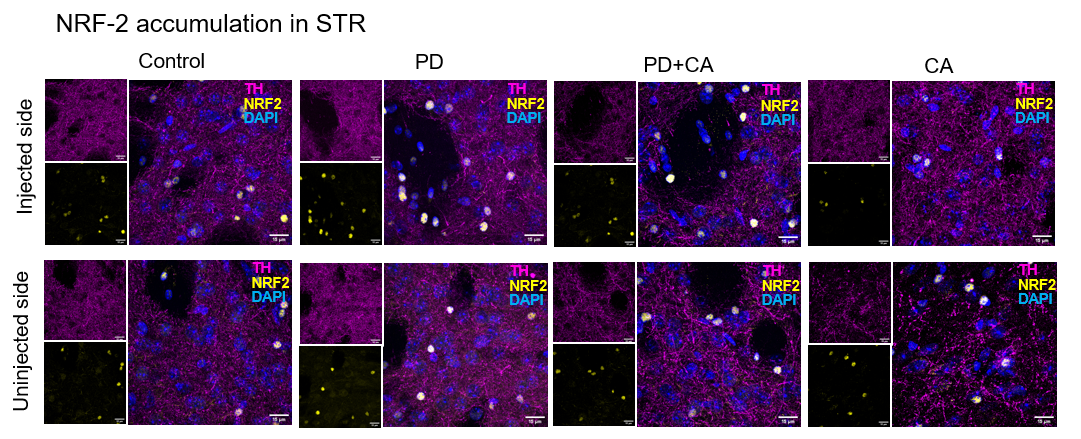

**Fig S6- NRF2 in the striatum (STR).** Representative images of STR co-stained with NRF2 (yellow), TH (magenta), and DAPI (blue) in the injected and uninjected sides across all groups. Scale bar: 15 μm.
